# Sphingosine-1-phosphate receptor 3 activation promotes sociability and regulates the expression of genes associated with anxiolytic-like behavior

**DOI:** 10.1101/2024.07.31.606030

**Authors:** Jose Castro-Vildosola, Chris-Ann Bryan, Nasira Tajamal, Sai Anusha Jonnalagadda, Akhila Kasturi, Jaqueline Tilly, Isabel Garcia, Renuka Kumar, Nathan T. Fried, Tamara Hala, Brian F. Corbett

## Abstract

We previously demonstrated that sphingosine-1-phosphate receptor 3 (S1PR3) in the medial prefrontal cortex (mPFC) prevents stress-mediated reductions in sociability. S1PR3 is a ubiquitously expressed G-protein coupled receptor that regulates immune system function, although its regulation of other biological processes is not well understood. Pharmacological activators of S1PR3 might provide important insights for understanding the neural substrates underlying sociability and/or serve as novel, preclinical treatments for social anxiety. Here we show that in mice, systemic injections of an S1PR3-specific agonist, CYM5541, promotes sociability in males and females whereas an S1PR3-specific antagonist, CAY10444, increases amygdala activation and promotes social anxiety-like behavior in females. S1PR3 expression is increased in the mPFC and dentate gyrus of females compared to males. RNA sequencing in the mPFC reveals that S1PR3 activation alters the expression of transcripts related to immune function, neurotransmission, transmembrane ion transport, and intracellular signaling. This work provides evidence that S1PR3 agonists, which have classically been used as immune modulators, might also be used as novel anxiolytics. S1PR3 might be an important hub gene for anxiolytic effects as it reduces inflammatory processes caused by stress and increases transcripts linked to anxiolytic neurotransmission.

**Highlights:**

- The Sphingosine-1-phosphate receptor 3 (S1PR3) agonist CYM5541 promotes sociability
- The S1PR3 antagonist CAY10444 reduces sociability and promotes anxiety-like behavior in females
- CAY10444 increases neuronal activity markers in the amygdala
- Pharmacological activation of S1PR3 regulates the expression of genes in the prefrontal cortex that control a wide range of biological processes, including increasing GABAergic neurotransmission and reducing inflammatory processes

## 1. Introduction

Social behavior is critical for survival and reproduction. In rodents, reduced social interaction is broadly considered social anxiety-like behavior as a wide range of stressors reduce social interaction^1–4^ and anxiolytic drugs increase social interaction^5^. While currently available anxiolytic drugs are effective at treating anxiety disorders^6^, they can also cause adverse effects like misuse and dependence^7^, cognitive and motor deficits^8–10^, and seizures when high-dose treatment is ended abruptly^11^. Identifying novel neural substrates underlying social anxiety-like behavior might lead to the development of novel therapeutics that can mitigate side effects more effectively and/or improve treatment strategies for certain individuals. Understanding neural substrates underlying social interaction in rodents will enhance our understanding of social behavior.

We previously used the resident-intruder paradigm of social defeat stress to identify sphingosine-1-phosphate receptor 3 (S1PR3), a ubiquitous G-protein coupled receptor, as a novel neural substrate that reduces social anxiety-like behavior in stressed male rats^3^. Resilient rats in this paradigm behave similarly to non-defeated controls, whereas vulnerable rats display social anxiety- and depression-like behavior^12,13^. We found that in male rats that are resilient to the adverse effects of stress, S1PR3 is increased in the medial prefrontal cortex (mPFC), a brain region that negatively regulates anxiety-like behavior^14^. We found that S1PR3 mRNA is reduced in the blood of veterans with posttraumatic stress disorder (PTSD) compared to combat-exposed veterans without PTSD and inversely correlated with PTSD symptom severity. We demonstrated that S1PR3 is increased in resilient rodents because glucocorticoid receptors (GRs), which are anti-inflammatory nuclear receptors activated by stress-induced corticosterone, are increased in the mPFC of resilient rats and increase S1PR3. S1PR3 mitigates inflammatory processes as S1PR3 knockdown in the mPFC increased microglia densities and the expression of tumor necrosis factors-alpha (TNFα) following stress. Mitigating inflammatory processes caused by stress provides an anxiolytic-like effect for S1PR3 as pharmacological inhibition of TNFα reversed the anxiogenic and pro-depressive effects caused by S1PR3 knockdown. S1PR3 also altered local field potentials in the mPFC and neuronal activity markers in the extended mPFC network, although the mechanisms were not known. We did not observe behavioral effects in non-stressed males, but we only manipulated S1PR3 in the mPFC^3^. The effects of systemically activating S1PR3 specifically have not been investigated.

FTY720 (fingolimod) is a non-specific S1PR modulator that treats multiple sclerosis (MS) by preventing lymphocyte egress via functional antagonism of S1PR1^15,16^. FTY720 is also a partial agonist for S1PR3^17^. In stressed mice^18^ and under baseline conditions^19^, FTY720 increases social interaction. In the brain, FTY720 inhibits histone deacetylases, prevents reductions in brain-derived neurotrophic factor (BDNF), and mitigates inflammatory gene expression^18–20^. FTY720 also improves social interaction in rodent models of autism^21,22^. In patients with MS, FTY720 reduces anxiety scores^23,24^, although confounding variables should be considered given the disease context.

Here we demonstrate that systemic treatment of mice with the S1PR3-specific agonist CYM5541 (CYM) increases social interaction whereas the S1PR3-specific antagonist CAY10444 (CAY) reduces social interaction in females and increases fear behavior. We show that S1PR3 expression is higher in multiple stress-related brain regions in females, which might explain why behavioral effects caused by S1PR3 antagonism are only observed in females. CAY-treated mice display increased neuronal activity markers in the amygdala. Using RNA-sequencing in PFC tissue, we found that CYM increases the transcription of a neuropeptide and enzymes linked to anxiolytic neurotransmission. Together, these findings demonstrate that S1PR3 increases social behavior and reduces social anxiety-like behavior. Our findings demonstrate that S1PR3, an important immune system modulator, is also important for neurotransmission.

## 2. Methods

### 2.1 Mice

10 to 14-week-old male and female C57BL/6 mice were used in this study. Littermates were matched between treatment groups. For tissue collection, mice were transcardially perfused with saline. Right hemibrains were frozen on dry ice for biochemistry. For immunohistochemistry, left hemibrains were postfixed in 4% paraformaldehyde (PFA) in phosphate buffered saline (PBS) for 48 hours followed by 72 hours in cryoprotectant (30% glycerol, 30% ethylene glycol in PBS). Mice are kept on a 12-12 light-dark cycle with lights being turned on at 7:00 and lights off at 19:00. All procedures were approved by the Rutgers University Institutional Animal Care and Use Committee.

### 2.2 Treatment with S1PR3 agonists and antagonists

CYM5541 (CYM) was purchased from Selleck Chemicals (cat. no. S6552). CAY10444 (CAY) was purchased from Cayman Chemical (cat. no. 10005033). On the same day as injection, CYM or CAY were dissolved in dimethyl sulfoxide (DMSO) to make a final solution of 1.25 mg/mL CYM or CAY in 5% DMSO in corn oil. Mice were treated with 10 mg/kg of CYM or CAY, a dose previously used for each drug in the absence of any noted side effects^25,26^. 5% DMSO in corn oil was used as control vehicle. No obvious signs of distress or immobility were observed.

### 2.3 Social interaction paradigm

Social interaction was performed 120 minutes following a second daily injection of CYM, CAY, or vehicle. Mice were placed in a 30 cm x 38 cm social interaction chamber with 9 cm x 9 cm corner zones. The social interaction zone is a 25.5 cm x 10 cm rectangle with an arc that reaches out to an additional 3.75 cm at its peak. The enclosure target mice are placed in is an upside down, metal pencil holder with perforated sides that is 7.5 cm in diameter. Treated mice are placed in the social interaction chamber for 150 seconds without the target mouse, removed for 30 seconds while a novel CD-1 target mouse is placed in its enclosure, and returned to the social interaction chamber for an additional 150 seconds. All behavior in the social interaction chamber without and with the target mouse is video recorded. Time in either corner and time in the social interaction chamber, excluding time on top of the target mouse enclosure, are used for analysis.

### 2.4 Immunohistochemistry

Immunohistochemistry was performed as previously described^3,27,28^. Cryoprotected brains were sectioned at 30 µm on a sliding microtome. Primary antibodies used were monoclonal mouse anti-GAD-67 (1:1000, EMD Millipore, MAB5406), rabbit anti-S1PR3 (1:300, Bioss, bs-7541R), guinea pig anti-NeuN (1:3000, EMD Millipore, ABN90) and rabbit anti-cFos (1:1000, Synaptic Systems, 226-008). Alexa Fluor™ goat anti-guinea pig 405 (Abcam, ab175678), goat anti-mouse 488 (Abcam, ab150113) and goat anti-rabbit 594 (Abcam, ab150080) were used as secondaries at a concentration of 1:200. Sections were blocked with 10% normal goat serum for one hour and incubated with primary antibodies overnight at 4°C. Sections were rinsed and incubated in secondary antibodies for 120 minutes. Sections were mounted and imaged on a Leica TCS SP8 confocal microscope. Immunoreactivity was determined by quantifying cell counts for c-Fos. S1PR3 levels were assessed by quantifying the mean pixel intensity of regions of interest using Fiji ImageJ. For measuring S1PR3 in the dentate gyrus, a region of interest was drawn around the dentate gyrus granule cell layer. S1PR3 was quantified in the mPFC as previously published^3^. For measuring S1PR3 in the prelimbic (PL) and infralimbic (IL) subregions of the mPFC, a 3 x 3 grid was overlaid on the PL or IL. Within each grid square, the pixel intensity of the three cells closest to the center were quantified. Naïve males and females were used to assess sex differences in S1PR3 expression.

### 2.5 RNA isolation, sequencing, and analysis

RNA extraction was extracted as previously described^19,27^. Prefrontal cortex samples from vehicle- and CYM-treated male and female mice were sub-dissected using a light microscope. Tissue was lysed using the Qiagen TissueLyser method and RNA was extracted using the Qiagen RNeasy Lipid Tissue Mini Kit (cat. No. 74804) per the manufacturer’s instruction. Samples were eluted using nuclease-free water and isolated RNA concentration and quality were assessed using a NanoDrop 2000 spectrophotometer. RNA libraries were prepared and sequencing was performed by the Rutgers University Genomics Center. mRNA was isolated by polyA selection and sequenced using the Illumina NovaSeq 6000 Sequencing System. The Rutgers Molecular and Genomics Informatics Core (MaGIC) assisted with analysis. FastQC was used to confirm RNA quality and trim sequence reads. Spliced Transcripts Alignment to a Reference (STAR) was used to align sequence reads to the mouse transcriptome^29^. Salmon was used to quantify the number of aligned sequence reads^30^. Differentially expressed genes (DEGs) were identified using DESeq2^31^. For each DEG, the Ensembl transcript identification number, the gene symbol, gene name, base mean, Log2 fold change (FC), FC, p-value, and adjusted p-value are provided in Supplementary Tables 1 (male), 2 (female), and 3 (male and female combined). DEGs with a fold change between 0.95 and 1.05 were removed analysis. Biological processes and molecular function for each DEG were identified using gene ontology (GO) enrichment analysis using Cytoscape 2.0^32,33^. All code used for RNA-seq analysis can be found here: https://github.com/jec423/RNA-seq/tree/main.

### 2.6 Statistics

Student’s t-test was used to identify differences between drug and control groups. Behavior and immunohistochemistry data were analyzed using GraphPad Prism 7. For each comparison, data that were three standard deviations from the mean were removed. One vehicle-treated male mouse was removed based on the amount of time spent in the social interaction zone without the target mouse present. Prior to identifying DEGs, outliers were identified using Principal Components Analysis and removed from subsequent analyses. DESeq2 was used to identify differentially expressed genes using a Benjamini-Hochberg method to correct for false discovery rate.

## 3. Results

### 3.1 S1PR3 agonism increases sociability in males and females

We hypothesized that systemic administration of CYM5541 (CYM, 10 mg/kg, i.p.), an S1PR3-specific agonist^25^, would increase social interaction two hours following an initial injection (Fig. 1A,B). In the absence of a target mouse, CYM-treated male mice spent similar amounts of time in the social interaction zone (Fig. 1C, p = 0.92, t_14_ = 0.099) and the corner (Fig. 1D, p = 0.64, t_14_ = 0.4843) compared to vehicle-treated controls. With the target mouse present, CYM-treated males spent more time in the interaction zone compared to vehicle-treated males (Fig. 1E, p = 0.029, t_14_ = 2.437), without any difference in time spent in corner (Fig. 1F, p = 0.74, t_14_ = 0.3379). In the absence of a target mouse, CYM-treated females spent similar amounts of time in the social interaction zone (Fig. 1G, p = 0.13, t_13_ = 1.599) and the corner (Fig. 1H, p = 0.82, t_13_ = 0.2366) compared to vehicle-treated controls. With the target mouse present, females spent more time in the interaction zone (Fig. 1I, p = 0.015, t_13_ = 2.806) and less time in the corner (Fig. 1J, p = 0.0043, t_13_ = 3.446) compared to vehicle-treated females. Together, these findings indicate that CYM increases sociability in males and females and reduces fear, as assessed by decreased time spent in corner, in females.

**Fig. 1.**
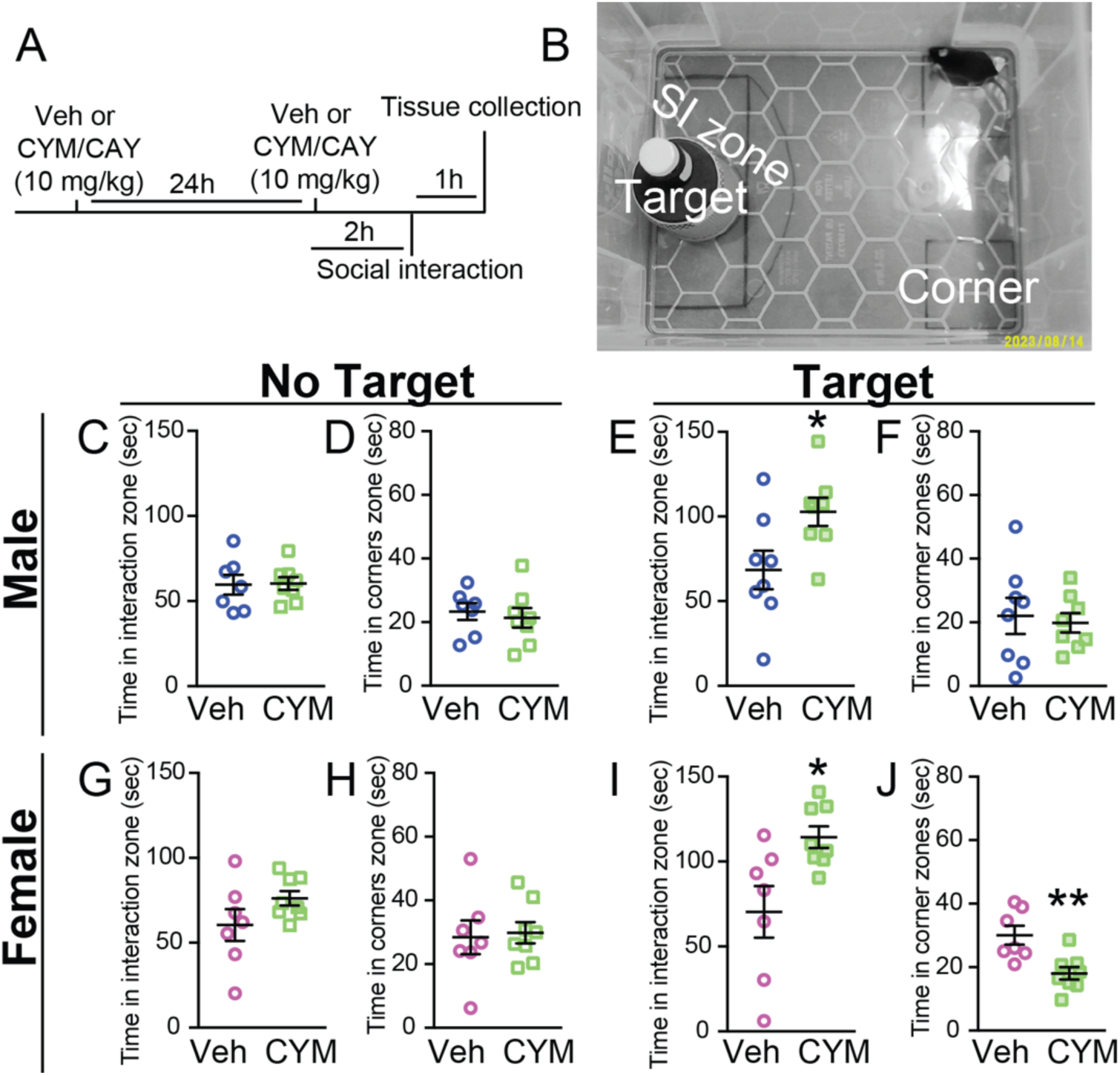
The S1PR3 agonist CYM5541 (CYM) increases sociability in males and females. A) Schematic for the timeline of treatment injections, social interaction assay, and tissue collection. Social interaction (SI) chamber with SI zone, corner, and target mouse zones labeled. Time in SI zone and D) corner in vehicle- and CYM-treated males without the target mouse present. Time in E) SI zone and F) corner in vehicle- and CYM-treated males with the target mouse present. Time in G) SI zone and H) corner in vehicle- and CYM-treated females without the target mouse present. Time in I) SI zone and J) corner in vehicle- and CYM-treated females with the target mouse present. For males, n = 8/group. For females, vehicle n = 7, CYM n = 8. Bars represent mean ± SEM. *p < 0.05, **p < 0.01. Veh vehicle; CYM CYM5541; CAY CAY10444; sec seconds.

### 3.2 S1PR3 antagonism increases social anxiety-like behavior in females, but not males

We hypothesized that the S1PR3-specific antagonist CAY10444 (CAY, 10 mg/kg, i.p.) would cause opposite effects of CYM. In the absence of the target mouse, CAY-treated males spent similar amounts of time in the social interaction zone (Fig. 2A, p = 0.2816, t_14_ = 1.12) and corner (Fig. 2B, p = 0.361, t_14_ = 0.994) compared to vehicle-treated males. With the target mouse present, CAY-treated males also spent similar amounts of time in the interaction zone (Fig. 2C, p = 0.7436, t_14_ = 0.3336) and corner (Fig. 2D, p = 0.9488, t_14_ = 0.065) compared to vehicle-treated males. In the absence of a target mouse, CAY-treated females spent less time in the interaction zone (Fig. 2E, p = 0.032, t_13_ = 2.406) and more time in the corner (Fig. 2F, p = 0.021, t_13_ = 2.638) compared to vehicle-treated females. CAY-treated females also spent less time in the interaction zone (Fig. 2G, p = 0.016, t_13_ = 2.755) and more time in the corner (Fig. 2H, p = 0.046, t_13_ = 2.198) compared to vehicle-treated females when the target mouse was present. These findings suggest that S1PR3 antagonism increases anxiety-like behavior in females, but not males, regardless of social interaction.

**Fig. 2.**
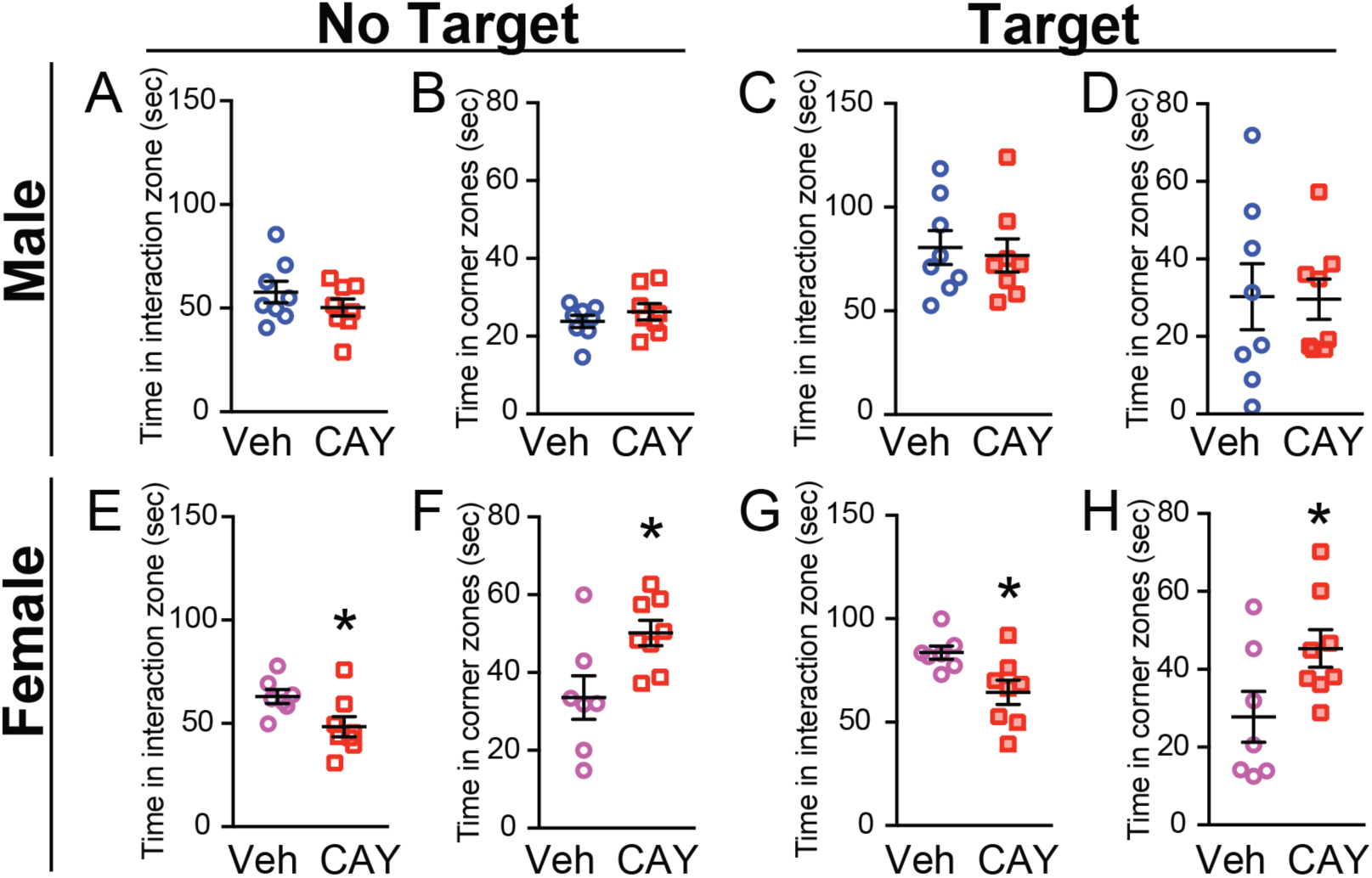
The S1PR3 antagonist CAY10444 (CAY) induces social anxiety-like behavior in females. Time in A) SI zone and B) corner in vehicle- and CAY-treated males without the target mouse present. Time in C) SI zone and D) corner in vehicle- and CAY-treated males with the target mouse present. Time in E) SI zone and F) corner in vehicle- and CAY-treated females without the target mouse present. Time in G) SI zone and H) corner in vehicle- and CAY-treated females with the target mouse present. For males, n = 8/group. For females, vehicle n = 7, CYM n = 8. Bars represent mean ± SEM. *p < 0.05. SI social interaction; Veh vehicle; CAY CAY10444; sec seconds.

### 3.3 S1PR3 in stress-related brain regions is higher in females compared to males

We hypothesized that S1PR3 expression might be higher in stress-related brain regions in females due to the stronger effects of S1PR3 antagonism. S1PR3 expression is higher in neurons compared to astrocytes and microglia, with similar levels in glutamatergic and gamma aminobutyric acid (GABA)ergic neurons^3^. S1PR3 expression has been reported in the hippocampus^34^ and mPFC^3^, which are both important for the behavioral and neuroendocrine responses to stress^3,14,35–39^. S1PR3 expression is higher in the dentate gyrus of naïve females compared to naïve males under baseline conditions (Fig. 3A-C, p = 0.0244, t_20_ = 2.434). S1PR3 is also higher in the prelimbic (PL, p = 0.0007, t_20_ = 3.941) and infralimbic (IL, p = 0.0022, t_20_ = 3.485) cortex subregions of the mPFC (Fig. 3, D-G) of naïve females compared to naïve males. These findings indicate that compared to males, female mice display increased expression of S1PR3 in the dentate gyrus and mPFC.

**Fig. 3.**
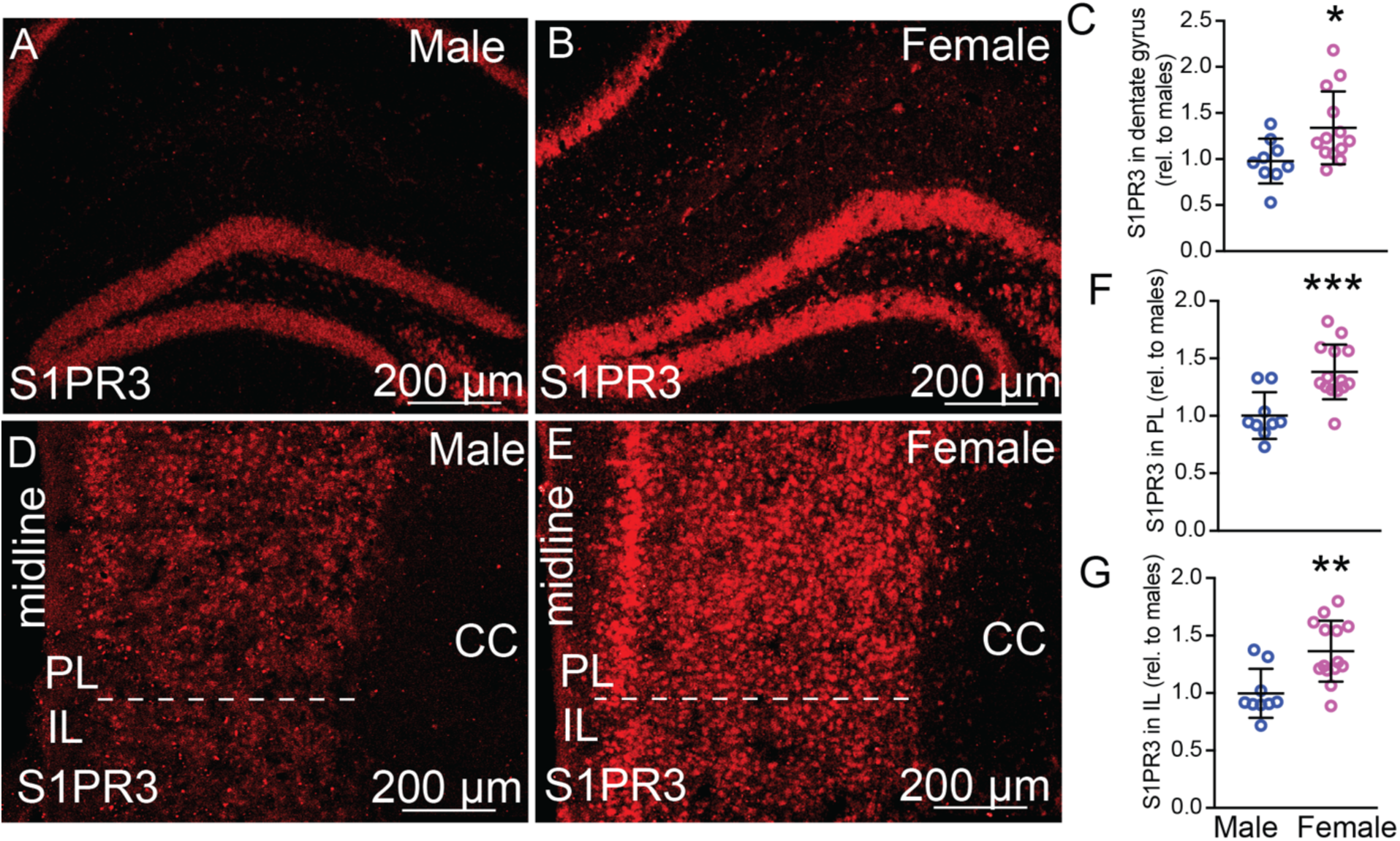
S1PR3 expression is higher in the brains of females compared to males. Images of S1PR3 in the dentate gyrus of A) male and B) female mice. C) Quantification of S1PR3 immunoreactivity in the granule cell layer of the dentate gyrus of naïve male and female mice. Images of S1PR3 in the mPFC of D) male and E) female mice. Quantification of S1PR3 in the F) PL and G) IL of male and female mice. For males, n = 9. For females, n = 13. Bars represent mean ± SEM. *p < 0.05, **p < 0.01, ***p< 0.001. CC corpus callosum; mPFC medial prefrontal cortex; PL prelimbic cortex; IL infralimbic cortex; rel relative.

### 3.4 S1PR3 antagonism increases neuronal activity markers in the amygdala

We and others have demonstrated that S1PR3 regulates neuronal activity^3,34,40,41^. We hypothesized that CAY treatment increases neuronal activity markers in the amygdala. Despite no changes in social interaction, CAY-treated males display a greater number of cFos-immunoreactive (IR) cells in the basolateral amygdala (BLA) (Fig. 4A-G, p = 0.019, t_13_ = 2.675) and central amygdala (CeA) (Fig. 4H, p = 0.011, t_13_ = 2.948) compared to vehicle-treated males. Compared to vehicle-treated females, CAY-treated females also display a greater number of c-Fos IR cells in the BLA (Fig. 4I, p = 0.018, t_13_ = 2.72) and CeA (Fig. 4J, p = 0.028, t_13_ = 2.436). Together, these findings demonstrate that S1PR3 is important for maintaining low levels of neuronal activity in the amygdala.

**Fig. 4.**
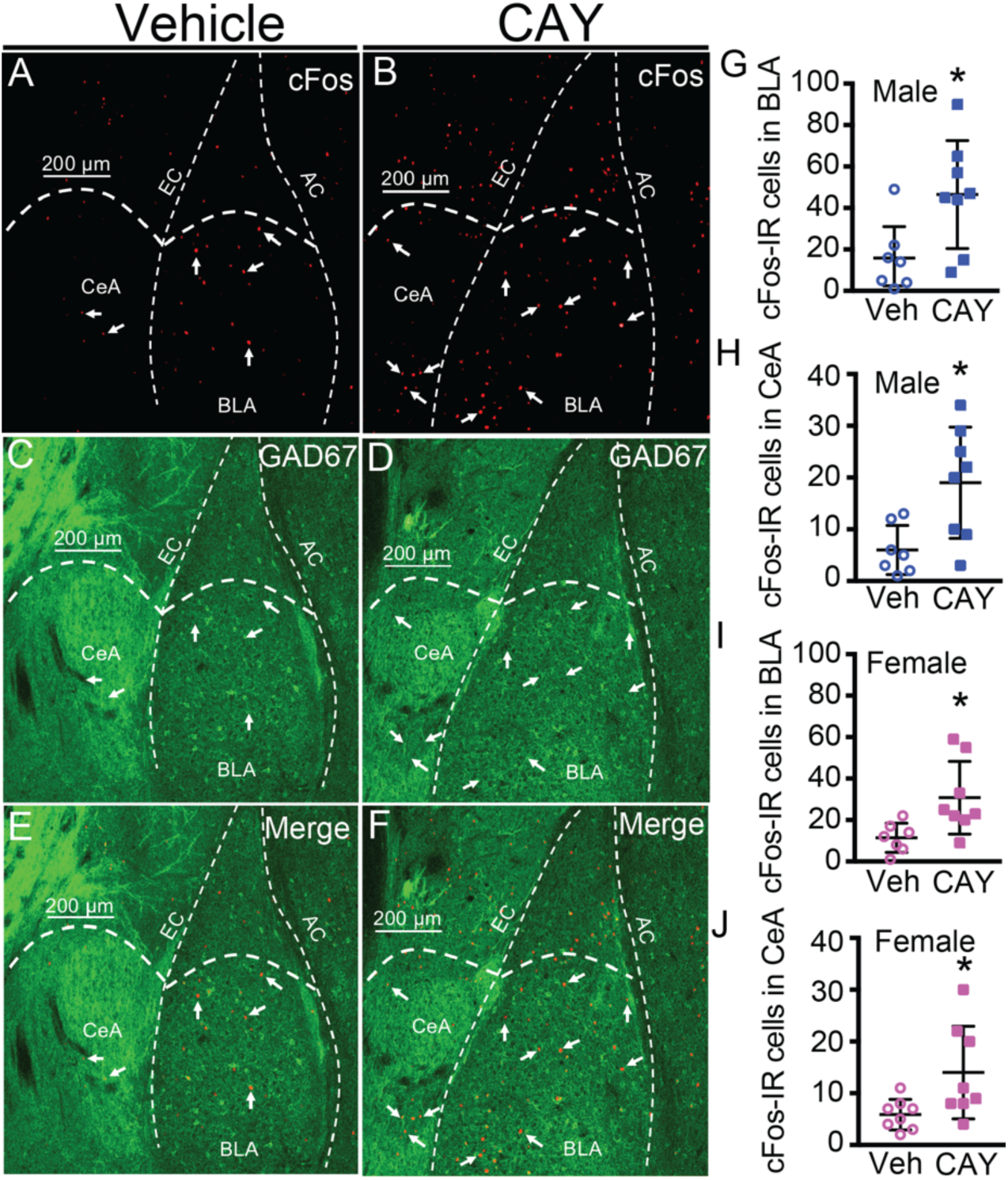
S1PR3 antagonism increases neuronal activity markers in the amygdala. A,B) c-Fos, C,D) GAD67, and E,F) merged images of the BLA and CeA of vehicle and CAY-treated males. Quantification of c-Fos in the G) BLA and H) CeA of veh- and CAY-treated males and I) BLA and J) CeA of veh- and CAY-treated females. Male veh, n= 7, male CAY n = 8, female veh n = 7, female CAY n= 8. Bars represent mean ± SEM. *p < 0.05. CeA central amygdala, BLA basolateral amygdala, EC external capsule, AC amygdalar capsule , veh vehicle, CAY CAY10444.

### 3.5 Transcripts in the prefrontal cortex altered by CYM

We performed RNA-sequencing (RNA-seq) on PFC tissue two hours following a second daily dose of CYM or vehicle. We chose the PFC because of our previous work demonstrating that S1PR3 in the PFC promotes resilience^3^. Compared to vehicle-treated males (n = 4), CYM-treated males (n = 5) displayed 11 transcripts that were decreased and 12 transcripts that were increased. Compared to vehicle-treated females (n = 5), CYM-treated females (n = 5) displayed 14 transcripts that were decreased and 43 transcripts that were increased. Combining males and females, CYM-treated mice displayed 125 transcripts that were decreased and 290 transcripts that were increased (Fig. 5A). All differentially expressed gene (DEG) abbreviations, full names, and statistics for males, females, and males and females combined are provided in Supplementary Tables 1-3.

**Fig. 5.**
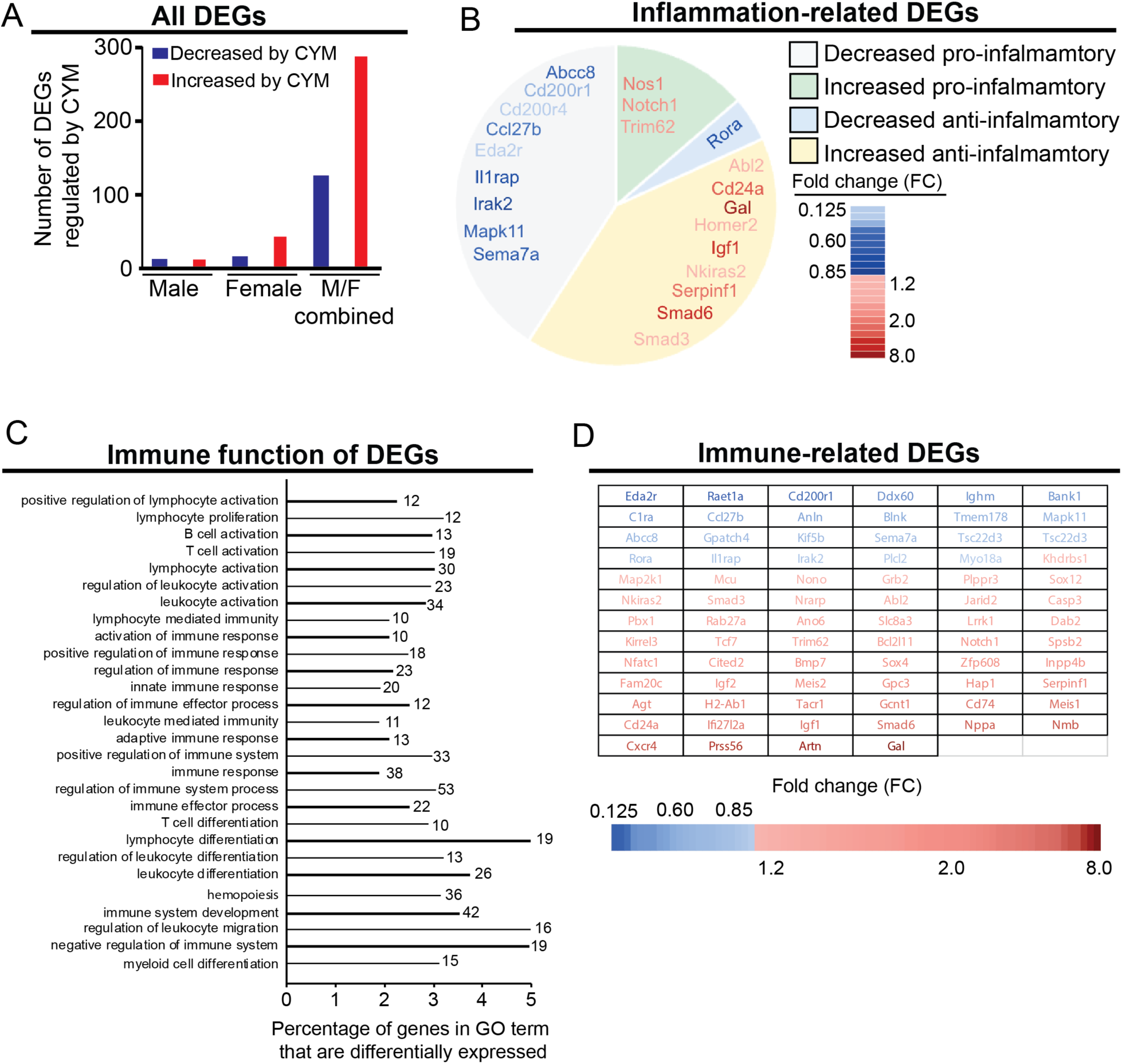
CYM reduces inflammation-related transcripts while increasing immune-related transcripts. A) Compared to vehicle-treated males (n = 4), CYM-treated males (n = 5) displayed 11 transcripts decreased and 12 increased DEGs. Compared to vehicle-treated females (n = 5), CYM-treated females (n = 5) displayed 14 decreased and 43 increased DEGs in the PFC. Combining males and females, CYM-treated mice displayed 125 decreased and 290 increased DEGs. B) Compared to veh-treated males and females, CYM increased nine anti-inflammatory and one pro-inflammatory transcript, and decreased nine pro-inflammatory and three anti-inflammatory transcripts in the PFC. Color is based on fold change. C) Percentage of genes associated with each immune-related GO term that are differentially expressed in the PFC of veh- and CYM-treated mice. The number after the bar indicates the number of DEGs. D) List of immune-related DEGs that are decreased or increased by in the PFC of CYM-treated males and female compared to veh-treated mice. Color is based on fold change. PFC prefrontal cortex, DEG differentially expressed gene, GO gene ontology, veh vehicle, CYM CYM5541.

We performed Gene Ontology (GO) enrichment analysis on vehicle and CYM-treated males and females to understand the molecular functions and biological processes regulated by S1PR3. Inflammatory transcripts were identified by their GO terms including “inflammation” as an associated biological process. Anti-inflammatory transcripts were associated with GO terms that had “negative regulation of inflammation” or “anti-inflammatory” as an associated biological process. Compared to vehicle-treated controls, 22 DEGs associated with inflammation were differentially expressed in the CYM-treated group. CYM decreased the expression of nine pro-inflammatory transcripts and one anti-inflammatory transcript. CYM increased the expression of three pro-inflammatory transcripts and nine anti-inflammatory transcripts (Fig. 5B). These DEGs, their fold change, and their adjusted p-values can also be found in Supplementary Table 4. These DEGs suggest an anti-inflammatory role of S1PR3 in the PFC under baseline conditions. This finding is consistent with our previous work demonstrating that S1PR3 mitigates stress-induced inflammation in the PFC^3^.

S1PR3 is important for regulating immune function in peripheral tissues, although the immune related genes regulated by S1PR3 in the brain are not well understood. We identified 22 transcripts related to immune function that are reduced by CYM and 53 immune-related transcripts that are increased by CYM. The GO terms associated with specific immune functions and the number of DEGs associated with each GO term were identified (Fig. 5C, Supplementary Table 5). These findings suggest that S1PR3 activation preserves immune function despite reducing baseline inflammatory transcription.

Mitigating inflammatory processes caused by stress increases sociability^3,42,43^. However, reducing inflammatory processes under baseline conditions, when inflammation is minimal, might not affect behavior. We hypothesized that activation of S1PR3 might increase sociability by increasing transcripts coding components of neurotransmitter systems that increase sociability. GABAergic agonists are widely prescribed as anxiolytics in humans^6^ and increase sociability in rodents^5^. We identified eight DEGs that are specifically related to GABAergic neurotransmission. Six of these were increased by CYM, including glutamate decarboxylase 1 and 2 (GAD 67 and GAD65, respectively), which are rate-limiting enzymes for GABA synthesis. Two GABA-related DEGs were decreased. Seven transcripts predicted to increase glutamatergic neurotransmission were increased in CYM-treated mice. The neuropeptide galanin, which exerts anxiolytic effects^44^ and reduces translation of TNFα^45^, was increased by CYM. Nitric oxide synthase 1 (Nos1), which promotes biosynthesis of nitric oxide (NO)^46^, was increased in CYM-treated mice. Angiotensin (Agt), which can reduce bioavailable NO^47^, and Smad3, which negatively regulates transcription of Nos^48^, were increased in CYM-treated mice. RAR-related orphan receptor alpha (Rora), which reduces reactive oxygen species^49^, was reduced in CYM-treated mice (Fig. 6A). 45 genes coding proteins directly involved in general or neurotransmitter-specific synaptic transmission were differentially expressed in CYM-treated mice compared to vehicle-treated controls (Supplementary Table 6).

**Fig. 6.**
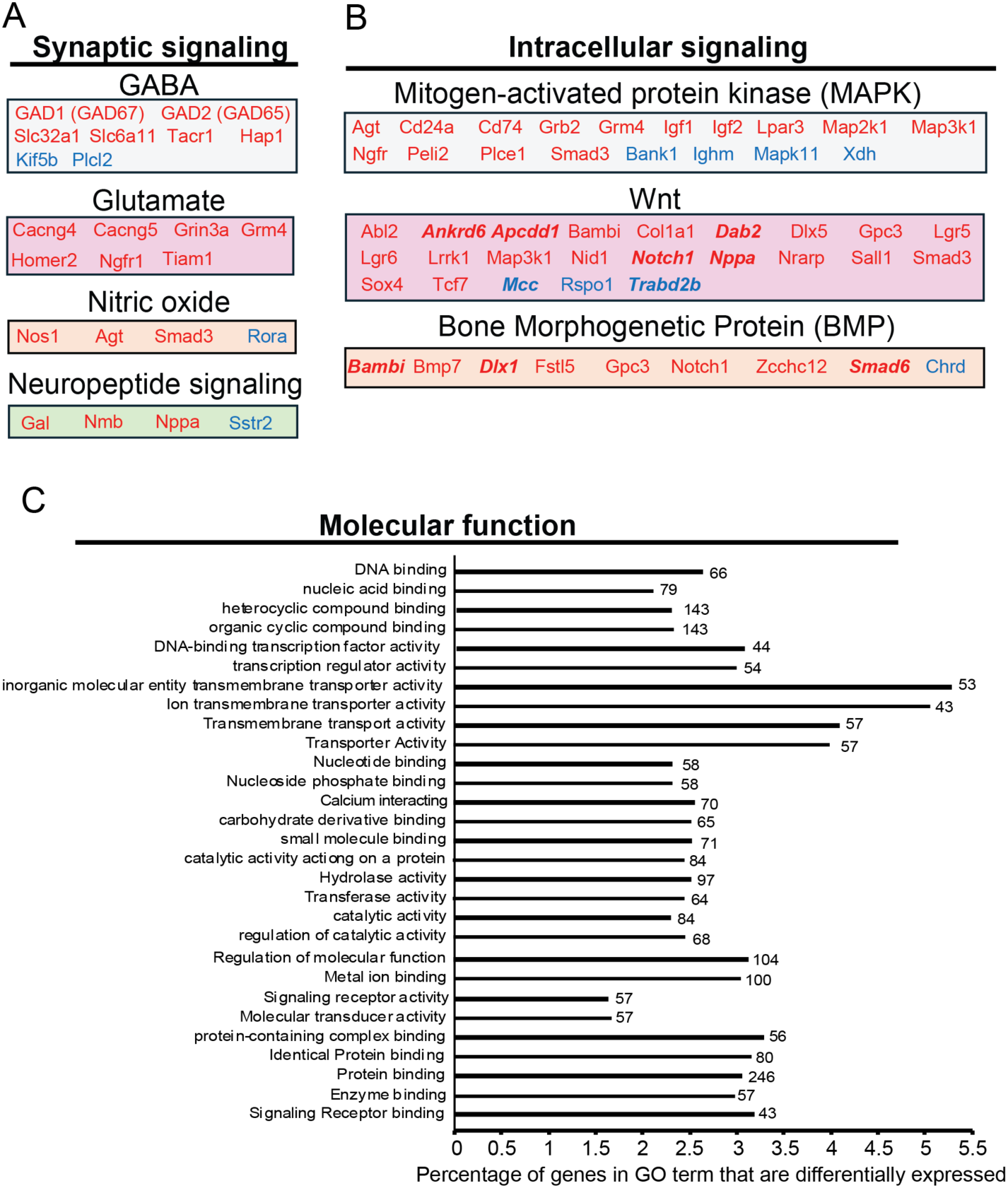
CYM regulates the expression of genes important for neurotransmission, intracellular signaling, and a wide range of other molecular functions. A) In the PFC, CYM-treated males and females display increased expression of six (red) and decreased expression of two (blue) genes that regulate GABAergic neurotransmission, increased expression of seven genes that regulate glutamatergic neurotransmission, increased expression of three and decreased expression of one gene that regulate nitric oxide neurotransmission, and increase three neuropeptides while decreasing one neuropeptide. B) In the PFC, CYM-treated males and females display differentially expressed genes that regulate MAPK, Wnt, and BMP signaling compared to veh-treated controls. Red represents DEGs with an increased fold change in CYM-treated males and females compared to veh-treated controls. Genes that are bolded and italicized have molecular function GO terms indicating they negatively regulate their intracellular signaling pathway. C) Percentage of genes associated with each molecular function GO term that are differentially expressed in the PFC of veh- (n = 9) and CYM (n = 10)-treated males and females. The number after the bar indicates the number of DEGs associated with that molecular function. PFC prefrontal cortex, DEG differentially expressed gene, GO gene ontology, veh vehicle, CYM CYM5541.

S1PR3 couples with multiple second messenger signaling pathways, including Gα_i_, Gα_q_, and Gα ^50^. These pathways, in turn, activate signaling cascades, like mitogen-activated protein kinase (MAPK) and calmodulin kinase (CaMK) pathways. Identifying changes in transcripts associated with specific pathways might represent enhanced function of those pathways. Four transcripts involved in MAPK signaling were reduced in CYM-treated mice and 14 were increased. Three transcripts associated with Wnt signaling pathways were decreased in CYM-treated mice. Two of these transcripts negatively regulate Wnt signaling, and one positively regulates Wnt signaling. Twenty transcripts associated with Wnt signaling were increased in CYM-treated mice. Only five of these up-regulated transcripts negatively regulate Wnt signaling. One transcript associated with bone morphogenetic protein (BMP) signaling was decreased in CYM-treated mice, and eight were increased. Three of these up-regulated transcripts negatively regulate BMP signaling (Figure 6B). For each signaling pathway, all DEGs are presented with their fold change and adjusted p-value in Supplementary Table 7. When possible, positive or negative regulation of each pathway is provided.

Molecular functions regulated by CYM and the number of DEGs associated with each function are provided (Fig. 6C, Supplementary Table 7). CYM-treated mice displayed a reduction in 13 transcripts that regulate transcription and an increase in 41 transcription-regulating transcripts. 44 of these transcriptional regulators directly bind DNA. Seven transcripts involved in transmembrane ion transport were decreased and 36 were increased in CYM-treated mice (Fig. 6C). We identified 70 genes that differentially regulated in CYM mice that bind calcium. 59 of these transcripts were increased, including CamK2d, and 11 were decreased. For each molecular function, all DEGs are presented with their fold change and adjusted p-value in Supplementary Table 7. Together, our findings demonstrate that S1PR3 promotes sociability, which might be attributed to altered expression of genes important for synaptic neurotransmission, intracellular signaling pathways, and/or immune function.

## 4. Discussion

Here we demonstrate that the S1PR3-specific agonist CYM increases sociability and reduces social anxiety in male and female mice. Conversely, the S1PR3-specific antagonist, CAY, reduces sociability and increases social anxiety-like behavior in females, but not males. This sex-specific effect might be attributed to S1PR3 levels being higher in females compared to males.

Consistent with increased social anxiety-like behavior in CAY-treated females, CAY increased neuronal activity markers in the BLA and CeA. Because we previously found that S1PR3 in the mPFC promotes stress resilience^3^, we performed RNA-seq using PFC tissue from CYM-treated males and female mice to identify the transcripts regulated by S1PR3. Consistent with higher levels of S1PR3 in females, we found that CYM alters the expression of more genes in females compared to males. Many of these genes are involved in immune function, inflammatory processes, neurotransmission, ion transport, intracellular signaling, and transcription. Together, these findings demonstrate that S1PR3, an immune-related G-protein coupled receptor, promotes sociability, reduces anxiety-like behavior, and regulates multiple biological processes that influence social behavior.

We previously found that S1PR3 prevents behavioral changes caused by stress by mitigating stress-induced tumor necrosis factor-alpha (TNFα) in the mPFC^3^. While immune and inflammation-related functions of S1PR3 has been extensively studied ^51–55^, S1PR3 controls other cellular processes, like neuronal activity^34,40,41^, which have not been studied as extensively. We can better understand these processes by assessing the effects of S1PR3 agonism under baseline conditions, in which inflammatory processes are minimal. In males, the S1PR3-specific agonist CYM increased sociability as assessed by increased time in the interaction zone with the target mouse present. Time spent in the interaction zone with the target mouse absent and time spent in the corner, regardless of whether the target mouse was present or absent, were not changed by CYM in males. This suggests that S1PR3 promotes sociability in males without affecting anxiety-like behavior as assessed by time spent in the corner. Time spent in the corner in vehicle-treated males is low, making reductions in this anxiety-like behavior difficult to detect under baseline conditions. Future studies will determine whether CYM reduces time spent in the corner in stressed males, which display increases in anxiety-like behavior^56^. Alternatively, the anxiolytic-like effects of CYM might be specific to social behavior in males. Similar to males, changes in behavior were not observed in CYM-treated females when the target mouse was absent, and CYM increased social interaction compared to vehicle-treated females. Unlike males, female CYM-treated mice spent less time in the corner when the target mouse was present, suggesting a stronger anxiolytic-like effect of CYM in females. Together, these findings indicate that pharmacological activation of S1PR3 increases sociability in males and females.

We found that in male mice, systemic injection of the S1PR3-specific antagonist, CAY, had no effect on time spent in the interaction zone or corner regardless of whether a target mouse was present. In females, CAY decreased time in the interaction zone and increased time in the corner regardless of the presence or absence of a target mouse. CAY-treated females might have spent less time in the interaction zone, even when the target mouse was absent, because they spent more time in the corner. This finding indicates that CAY does not affect anxiety-like behavior in males but increases anxiety-like behavior in females. Therefore, baseline S1PR3 activation is required to prevent anxiety-like behavior in females. Identifying mechanisms underlying anxiety-like behavior in females might have translational potential as stress-related mood disorders are twice as prevalent in women compared to men^57^.

We hypothesized that the stronger effects of CAY in females are attributed to higher levels of S1PR3. We found that S1PR3 is increased in the mPFC and dentate gyrus of females compared to males. We investigated the mPFC since we had previously studied the expression and function of S1PR3 in that region^3^. We investigated the dentate gyrus because S1PR3 regulates hippocampal neuron excitability^34^. We previously found that S1PR3 is increased by glucocorticoid receptors (GRs)^3^, which are expressed in the mPFC^3,58^ and dentate gyrus^59^. We hypothesize that S1PR3 is higher in females compared to males because baseline levels of the endogenous GR agonist, corticosterone, are higher in females^60^. To the best of our knowledge, a comprehensive brain-wide mapping of S1PR3 protein has not been performed. Therefore, S1PR3 might be increased in other brain regions in female mice compared to males. Increased expression or function of S1PR3 in females could have translational potential as stress-related disorders are twice as prevalent in women compared to men^57^.

We found that S1PR3 antagonism with CAY increases counts of the neuronal activity marker, c-Fos, in the BLA and CeA. This is consistent with increased time spent in the corner in CAY-treated females and supports the hypothesis that S1PR3 lowers anxiety-like behavior under baseline conditions. There are likely multiple mechanisms by which S1PR3 maintains low levels of neuronal activity in the amygdala. We previously found that S1PR3 overexpression in the mPFC reduces c-Fos in the BLA and CeA of stressed males^3^. The PFC reduces amygdala output via glutamatergic projections to inhibitory interneurons in the BLA^61^. S1PR3 might increase baseline activity of these neurons to inhibit the amygdala. However, S1PR3 might reduce the overall activity of the PFC because our RNA-seq experiment found that CYM reduces transcription of activity regulated cytoskeleton-associated protein (Arc), a neuronal activity marker^39,62^, in females. Others groups have found that S1PR3 promotes neuronal excitability in the hippocampus^34^ and periphery by activating transient receptor potential channels^40^ and closing KCNQ2/3 channels^41^. In the mPFC, S1PR3 is expressed at similar levels in glutamatergic and GABAergic neurons^3^. The effects of S1PR3 on neuronal activity likely vary across brain regions and cell type. We hypothesize that a combination of factors regulated by S1PR3 that influence neuronal activity in the amygdala and its afferents reduce amygdala activation under baseline conditions. Our RNA-seq data provide evidence that these factors also include altered expression of genes important for synaptic neurotransmission.

We found that CYM increases the expression of six genes in the PFC that are associated with enhanced GABAergic neurotransmission. Among these are GAD-67 and GAD-65, rate-limiting enzymes in GABA production^63,64^. CYM-treated mice also displayed increases in inhibitory synaptic factor family member 2b (Insyn2b), which inhibits post-synaptic potentials^65^. Increasing GABAergic neurotransmission might contribute to increased sociability^5^. However, the regulation of synaptic transmission by S1PR3 activation is not strictly inhibitory. S1PR3 also increased the expression of seven genes linked to glutamatergic neurotransmission. CYM-treated mice displayed increased transcription of galanin, which exerts anxiolytic-like effects when binding to the Gal-R1 and R2 receptors^44,66^, but promotes anxiety when binding to Gal-R3^67^. Galanin also reduces the translation of TNFα^45^. This provides a potential mechanism for our previous finding that S1PR3 reduces TNFα in the mPFC^3^. S1PR3 increased the expression of genes linked to the synthesis of NO, a retrograde neurotransmitter. NO controls anti-inflammatory effects under normal conditions^68^, but induces an inflammatory response when produced in excess^69^. This might at least partially explain the opposing effects of S1PR3 on inflammatory processes in different disease models. S1PR3 reduces inflammatory processes in rodent models with lower NO levels, like psychological stress^3^, obesity^53^, and baseline conditions^70^, but increases inflammatory processes in mouse models associated with excess NO levels like cancer^71^, lipopolysaccharide injection^72^, and stroke^73,74^. Indeed, the pro-inflammatory of S1PR3 in stroke require NO^74^. We hypothesize that S1PR3 reduces inflammatory processes by synthesizing NO synthesis at low levels. However, when NO levels are elevated by pro-inflammatory conditions, S1PR3 contributes to inflammatory processes by augmenting NO levels to excess and by priming immune function.

Our GO analysis found that 53 of the genes increased by CYM are associated with immune function. S1PR3 might play a role in priming the immune system to enhance innate and adaptive immunity. DEGs regulated by CYM were associated with the differentiation, development, and migration of myeloid cells, T cells, and B cells. Priming the immune system could lead to increased inflammatory processes following infection or tissue damage. The source of these transcripts is likely parenchymal brain cells, including microglia, because blood immune cells were mostly eliminated by transcranial perfusion. Future studies will investigate the role of S1PR3 in specific immune cell types.

S1PR3 regulates a wide range of biological processes, which might be attributed to its regulation of multiple intracellular signaling pathways. We found that CYM altered the expression of the expression of genes involved in MAPK, Wnt, and BMP signaling, along with 70 genes that interact with calcium. Altered expression of CamK2D and the other 69 transcripts encoding calcium-binding proteins might reflect increases in calcium release from the endoplasmic reticulum caused by Gα_q_ signaling, which is regulated by S1PR3^50^. The expression of calcium-binding proteins might be altered to enhance calcium signaling or compensate for increased cytosolic calcium concentrations^75^. Parvalbumin, a calcium buffer expressed by a subset of GABAergic interneurons^76^, was reduced in the PFC of CYM-treated mice. This might promote sociability and reduce anxiety-like behavior, especially in females, because chronic stress increases parvalbumin in the PFC, which contributes to anxiety-like behavior in females^77^. S1PR3 regulates MAPK signaling^73^, but to the best of our knowledge, has not been implicated in CaMK, Wnt, or BMP signaling. However, other S1PRs have been shown to regulate these pathways^78–81^. Therefore, S1PR3 might regulate these pathways or regulate the expression of components of these pathways to allow for cross-talk among S1PRs.

One caveat of this work is that CYM and/or CAY might cause off-target effects that regulate biological processes independent of S1PR3^20^. However, similar effects on sociability caused by S1PR3 overexpression^3^ or pharmacological activation of S1PR3 by different drugs^82^ minimizes this possibility. A caveat of GO analysis is that all the biological processes regulated by a gene cannot be completely known and might vary across cell types. Regardless, many of the biological processes confirmed by GO analysis have been confirmed by literature searches and background knowledge. Further, results are generally corroborated by identifying multiple DEGs with similar functions. Therefore, GO analysis provides important insights into complex changes in the transcriptome and how they might regulate biological processes. A third caveat is that changes in gene expression are correlative. DEGs might respond to CYM to enhance S1PR3 signaling or compensate for changes caused by it. We care able to hypothesize whether DEGs enhance S1PR3 signaling or compensate for it by understanding the biological processes regulated by S1PR3. Future work will investigate this directly by determining whether specific biological processes regulated by CYM are dependent on specific DEGs. An important caveat into the translational potential of using S1PR3 agonists is that S1PR3 promotes expansion of tumor cells via Notch signaling and MAPK^83–85^. Indeed, we show here that CYM increases transcription of genes in these pathways. Inhibiting MAPK and Notch might allow CYM treatments to be administered with fewer adverse side effects.

Together, our findings demonstrate that S1PR3 promotes sociability and prevents anxiety-like behavior in females. We demonstrate that S1PR3 is important for reducing neuronal activity markers in the amygdala. We found that S1PR3 regulates the transcription of genes associated with a wide range of biological functions, including reducing inflammatory processes and regulating neurotransmission. Of note, pharmacological S1PR3 activation increased the expression of GAD-67 and GAD-65, the rate-limiting enzymes in GABA production. S1PR3 is well-positioned for regulating stress-related behaviors as its expression is increased by GRs, it is reduced in veterans with PTSD, and it regulates multiple biological processes that reduce maladaptive behaviors caused by stress.

## 5. Conclusions

These findings indicate that S1PR3 is an important regulator of social behavior, especially in females. S1PR3 regulates the expression of genes that control a wide range of molecular functions linked to anxiolytic behavior, including genes related to GABAergic neurotransmission and reducing inflammatory processes.

## CRediT authorship statement

**Jose Castro-Vildosola:** Methodology, Software, Formal analysis, Investigation, Data curation, Writing – original draft, Writing -review and editing, Visualization. **Chris-Ann Bryan:** Methodology, Investigation, Writing -review and editing. **Nasira Tajamal:** Investigation. **Sai Anusha Jonnalagadda:** Methodology, Investigation, Writing -review and editing. **Akhila Katsuri:** Methodology, Investigation, Writing -review and editing. **Jaqueline Tilly:** Methodology, Investigation. **Isabel Garcia:** Methodology, Investigation. **Renuka Kumar:** Methodology, Investigation. **Nathan T. Fried:** Supervision. **Tamara Hala:** Methodology, Formal analysis, Investigation, Data curation, Writing – original draft, Writing -review and editing, Visualization, Supervision, Project administration. **Brian F. Corbett:** Conceptualization, Methodology, Software, Formal analysis, Investigation, Data curation, Writing – original draft, Writing -review and editing, Visualization, Supervision, Project administration, Funding acquisition.

## Acknowledgement and funding sources

We would like to thank Dr. Mainul Hoque, Dr. Patricia Soteropoulos, Andreca Frater, and Dr. Veera D’Mello at the Rutgers New Jersey Medical School Genomics Center for their assistance with RNA sequencing. We would like to thank Dr. Alexander Lemenze at the Rutgers New Jersey Medical School Molecular and Genomics Informatics Core for his assistance with the analysis with RNA sequencing data. This project was funded using start-up funds awarded to BC from the Faculty of Arts & Sciences at the Camden campus of Rutgers, The State University of New Jersey.

